# Association of ocean macroplastic debris with stranded sea turtles in the Central Gulf of Thailand

**DOI:** 10.1101/2021.10.15.464521

**Authors:** Jindarha Prampramote, Worakan Boonhoh, Sutsiree Intongead, Watchara Sakornwimol, Pimchanok Prachamkhai, Chalutwan Sansamur, Orachun Hayakijkosol, Tuempong Wongtawan

## Abstract

The impact of macroplastic debris (> 5 mm in size) on marine life is a global concern but is rarely investigated in Thailand. Our objective was to investigate the relationship between stranded sea turtles and macroplastics found in the Central Gulf of Thailand. The turtle (n = 388) stranding record between 2017-2020 was analysed retrospectively to determine their size, species, and interaction with macroplastics. Thereafter, between 2019-2020, macroplastics were collected, from the gastrointestinal (GI) tract of dead turtles and from the beaches where stranded turtles were found. A stereomicroscope was used to visually categorise the macroplastics, and the plastic composition was analysed using a Fourier-transform infrared (FTIR) spectrometer. Green sea turtles (*Chelonia mydas*) were found to account for the majority of stranded turtles (74%, n = 251), and macroplastics were discovered in 74% of cases of entanglement and ingestion. At the juvenile stage, the stranded turtle was strongly related to macroplastics. Immature turtles were more likely to become entangled than adult turtles. Entangled turtles had a greater survival rate than turtles that had consumed plastic. The plastic fibres were the majority of macroplastics found in the GI tracts (62%, n = 152/244) and on the beach (64%, n = 74/115). Most fibres from GI tracts (83%, n = 126/152) and the beaches (93%, n = 68/74) were identified as the fishing net comprised of polyethylene or polypropylene. We concluded that fishing nets made of polyethylene or polypropylene might be one of the significant causes of sea turtle stranding in the Central Gulf of Thailand, and this issue requires immediate resolution.

## 1. INTRODUCTION

Ocean plastic debris is a global issue, impacting over 900 marine species through ingestion and entanglement (Ryan 2016, Reinert et al. 2017, Kuhn & Van Franeker 2020). Depending on its size, marine plastic debris is classified as macroplastics (> 5 mm), microplastics (1 µm - 5 mm), or nanoplastics (< 1 µm) (Merga et al. 2020). While micro- and nanoplastics can be directly consumed by small organisms and accumulate in the food web (Diepens & Koelmans, 2018), macroplastics pose a particular problem for large marine animals such as manatees, whales, and turtles as well as sea birds, which become entangled in or ingest large debris such as fishing nets, potentially resulting in interruption and obstruction of the gastrointestinal (GI) tract (Jacobsen et al. 2010, Reinert et al. 2017, Duncan et al. 2019, Baak et al. 2020). Not only can macroplastics clog the GI tract, but they can also be harmful to animals that consume them due to the persistence of certain toxic chemicals such as plasticisers (Lithner et al. 2011). Additionally, toxic substances (e.g., heavy metals, polycyclic aromatic hydrocarbons, and polychlorinated biphenyls) can accumulate in plastics in the ocean (Mato et al. 2001, Nakashima et al. 2012, Bouhroum et al. 2019) and certain pathogenic microbes (e.g., vibrio and pseudomonas) can adhere to the surface layer of marine plastics (Kirstein et al. 2016, Viršek et al. 2017, Wu et al. 2019). These chemicals and pathogens have a potential to cause serious diseases in marine animals and jeopardise food security and food safety of human (Derraik, 2002, Teuten et al., 2007, Brennecke et al. 2016, Barbora et al. 2018).

Thailand is the world’s top ten producers of marine plastic (Jambeck et al. 2015). Nonetheless, research on plastic debris in the Thai ocean (the Andaman Sea and the Gulf of Thailand) has been limited to two studies that determined the incidence of microplastics in demersal and pelagic fishes (Azad et al. 2018, Klangnurak & Chunniyom 2020). According to local media and government agencies, macroplastics are occasionally discovered in the digestive tracts of stranded sea turtles (https://www.bangkokpost.com). Four species of sea turtles are recently reported in the Gulf of Thailand and the Andaman sea by the Department of Marine and Coastal Resource (https://www.dmcr.go.th); the majority population are green sea turtles *(Chelonia mydas)*, followed by hawksbill sea turtles (*Eretmochelys imbricata*), while olive ridley sea turtles (*Lepidochelys olivacea*) are uncommon, and leatherback sea turtles *(Dermochelys coriacea)* are extremely rare. According to the International Union for Conservation of Nature (IUCN, https://www.iucnredlist.org) report that in the Asian ocean, green sea turtles are categorised as endangered species, the hawkbills sea turtles are critically endangered, while olive ridley sea turtles and leatherback sea turtles are vulnerable.

According to the Department of Marine and Coastal Resource (https://www.dmcr.go.th), sea turtle population and nests in Thailand are decreasing each year which may be attributed to tourism, fishing, limited nesting area, and pollutions. However, the causes for the population decline and stranding have not been extensively studied. For the present study, we hypothesised that macroplastics were one of the leading causes for the stranding and death of sea turtles in the Central Gulf of Thailand. The objective of this study was to investigate the association of macroplastics and the stranding of sea turtles in the Central Gulf of Thailand.

## 2. MATERIALS AND METHODS

This study was approved by the Institutional Animal Care and Use Committee (IACUC) of Walailak University (project number 63-009).

### 2.1. Data of stranded sea turtle

To determine the relationship between marine plastics and stranding of sea turtles, firstly, we performed a retrospective analysis of the record of 338 stranded turtles along the shores of the Central Gulf of Thailand in 3 provinces (Chumphon, Surat Thani, and Nakhon Si Thammarat; Fig. 1) between 1 January 2017 and 31 July 2020. The stranded turtles were found and reported by local residents and fishermen. Thereafter, staff from the Marine Animal Research and Rescue Centre of Walailak University or the Marine and Coastal Resources Research Centre went to retrieve the turtles in order to investigate the cause of stranding and health status or cause of death according to standard protocols (Work 2000, Flint et al. 2009, Werneck et al. 2018). For dead turtles, the carcass condition was categorised to fresh, evident decomposition, advanced decomposition, or mummified (Werneck et al. 2018).

**Fig. 1.**
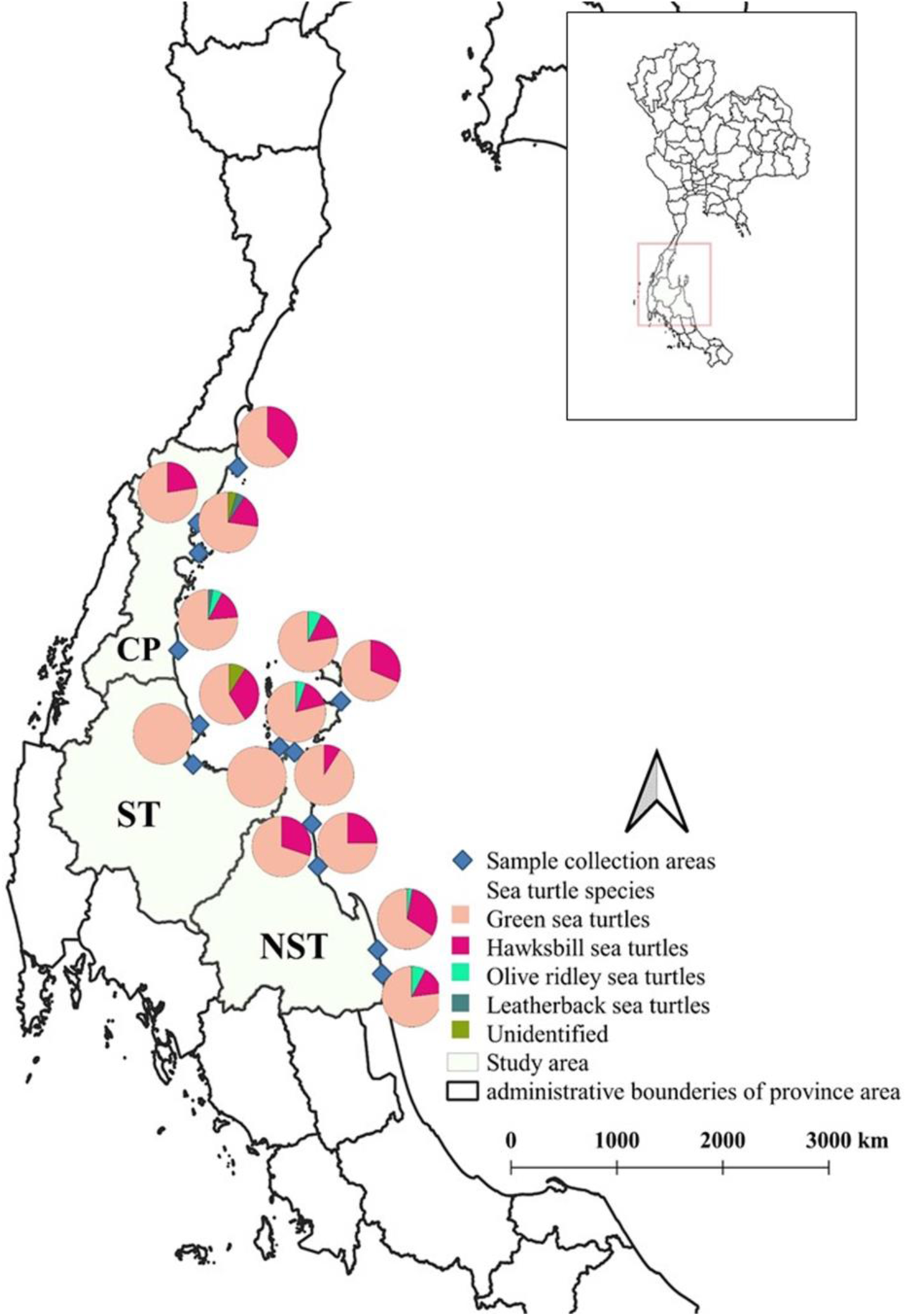
Study locations at the Central Gulf of Thailand included Chumphon, Surat Thani, and Nakhon Si Thammarat provinces. Black triangles represent the sample collection areas. Pie charts represent frequencies of differences stranded sea turtle species. Blue spots represent sample collection areas.

The sexes of deceased turtles were determined during necropsy or by examining the tail morphology of live adult turtles (adult males have a significantly longer tail than females), but live juveniles and sub-adults could not be sexed (Schofield et al. 2017). Each stranded turtle was grouped according to its size based on previous studies (Bresette et al. 2010, Jensen et al. 2018, Robinson et al. 2021) in the following categories. Green sea turtles, a juvenile (< 65 cm curved carapace length [CCL]), sub-adult (65-86 cm CCL), and adult-sized (> 86 cm CCL). Hawksbill sea turtles, juvenile (<55 cm CCL), sub-adult (>55–70 cm CCL) or adult (>70cm CCL). Olive ridley sea turtles, juvenile (<62 cm CCL), sub-adult (62–70 cm CCL) or adult (>71 cm CCL). Leatherback sea turtle, juvenile (<50 cm CCL), sub-adult (50– 70 cm CCL) or adult (>70 cm CCL).

### 2.2. Plastic analysis

After performing preliminary retrospective analysis, we discovered the evidence that macroplastics may be related to sea turtle stranding. Additionally, the data collected from the Marine and Coastal Resources Research Centre (https://km.dmcr.go.th/en/c_6/d_982) reports that green sea turtles in the Gulf of Thailand usually do not move far from the shore As a result, we would like to further investigate the type of plastics and the association of plastics found in the GI tract of turtles and on the beach. We hypothesised that plastics found on the beaches might be similar to the feeding area of marine turtles in the ocean.

Therefore, macroplastics were collected from the turtles and the beaches between 2019 and 2020 for further analysis. There were 244 pieces of macroplastics obtained from the stomach and small intestine of 30 dead stranded turtles (only fresh and evident decomposition stage).

On the beach, ocean macroplastics (size > 5 mm) were collected along a 100-metre transect mid-way between high and low tide (shoreline width < 6 m) following the sampling procedure by a previous study (Besley et al. 2017) on 13 beaches where stranded sea turtles were commonly observed (Fig. 1). For Chumphon province, macroplastics were collected from Sairee beach of Muang district, Hat Kho Khao of Lang Suan district, the Fishing village of Thung Tako district, Thung Wua Laen Beach of Pathio district, and Tongsai of Sawi district. For Surat Thani province, macroplastics were collected from Ferry Terminal of Donsak district, Laem Sai Beach of Chaiya district, and Leeled Beach of Punpin District. For Nakhon Si Thammarat province, macroplastics were collected from Khwaeng Phao Beach of Khanom District, Bang Dee Beach of Sichon District, Ban Tha Sung Bon Beach of Thasala District, Koh Fai Beach of Pak Phanang District, and Chan Chaeng Beach of Hausai. Totally, 155 macroplastics were collected from beaches.

Macroplastics were visually categorised as fibre, bag, foam, straw (for drinking), or hard plastic, followed by confirmation under a stereomicroscope (SMZ460 Zoom, Nikon Instruments) using reference plastic materials obtained from local fishermen and stores. A Fourier-transform infrared (FTIR) spectrometer (Tensor 27 equipped with a Platinum-ATR-unit; Bruker Optik) was used to determine the chemical composition of macroplastics. We co-added 32 scans to achieve an appropriate signal-to-noise ratio, with a spectral resolution of 8 cm^−1^ (per centimetres) in a wavenumber ranges from 4000 to 400 cm^−1^ (Primpke et al. 2018). Obtained spectra were analysed with the software OPUS 7.5 (Bruker Optik) through comparison with polymer reference spectra from an in-house plastic database and a previous study (Jung et al. 2018a).

### 2.4. Statistical analysis

Binomial regression was used to predict the probability of cause of stranding and macroplastics problem including (entanglement in fishing nets or ingestion leading to gastrointestinal obstruction) based on the descriptive variable species (Green sea turtles, Hawksbill sea turtles and Olive ridley sea turtles) and life history stages (juvenile, sub-adult and adult). A chi-square analysis to determine differences between entanglement and ingestion probabilities for the death of sea turtles. All analyses were performed using R statistical software version 4.0.4.

## 3. RESULTS

### 3.1. Occurrence of stranded turtles

Between 2017 and 2020, 338 stranded turtles (130 live and 208 dead) were found along the shores of the Central Gulf of Thailand. The majority of stranded turtles were green sea turtles (*Chelonia mydas*) (74.26%, n = 251) and hawksbill sea turtles (*Eretmochelys imbricata*) (21.01%, n=71). Others were olive ridley sea turtles (*Lepidochelys olivacea*) (3.25%, n =11), Leatherback sea turtles (*Dermochelys coriacea*) (0.59%, n = 2). However, several carcasses were unable to be identified, resulting from severe damage (0.88%, n = 3) (Table 1).

**Table 1.** Stranding information and cause of stranding for different turtle species in the Central Gulf of Thailand.

| Species | % of stranded turtles<br>(n) |  |  |  |  | % Causes of stranding<br>(n) |  |  |  |
| --- | --- | --- | --- | --- | --- | --- | --- | --- | --- |
|  | Viability |  | Stage of life |  |  | Macroplastics<sup>2</sup> | Health problem<sup>3</sup> | Injury<sup>4</sup> | Unknown |
|  | Live | Dead | Juvenile | Sub-adult | Adult |  |  |  |  |
| Green sea turtles<br>(n = 251) | 28.69%<br>(72) | 71.31%<br>(179) | 60.96%<br>(153) | 21.91%<br>(55) | 17.13%<br>(43) | 43.03%<br>(108) | 9.96%<br>(25) | 3.98%<br>(10) | 43.02%<br>(108) |
| Hawksbill sea turtles<br>(n = 71) | 78.87%<br>(56) | 21.13%<br>(15) | 81.69%<br>(58) | 12.68%<br>(9) | 5.63%<br>(4) | 59.15%<br>(42) | 21.13%<br>(15) | 1.41%<br>(1) | 18.31%<br>(13) |
| Olive ridley sea turtles<br>(n = 11) | 18.18%<br>(2) | 81.82%<br>(9) | 45.45%<br>(5) | 36.36%<br>(4) | 18.18%<br>(2) | 18.18%<br>(2) | 0.00%<br>(0) | 0.00%<br>(0) | 81.82%<br>(9) |
| Leatherback sea turtles<br>(n = 2) | 0.00%<br>(0) | 100.00%<br>(2) | 100.00%<br>(2) | 0.00%<br>(0) | 0.00%<br>(0) | 0.00%<br>(0) | 0.00%<br>(0) | 0.00%<br>(0) | 100.00%<br>(2) |
| Unidentified species<sup>1</sup> | 0.00%<br>(0) | 100.00%<br>(3) | 100.00%<br>(3) | 0.00%<br>(0) | 0.00%<br>(0) | 0.00%<br>(0) | 0.00%<br>(0) | 0.00%<br>(0) | 100.00%<br>(3) |
| **Total** | **38.46%**<br>**(130)** | **61.54%**<br>**(208)** | **65.38%**<br>**(221)** | **20.12%**<br>**(68)** | **14.50%**<br>**(49)** | **44.97%**<br>**(152)** | **11.83%**<br>**(40)** | **3.25%**<br>**(11)** | **39.94%**<br>**(135)** |

<sup>1</sup>Badly damaged cadavers (mummified), <sup>2</sup>Entanglement and/or ingestion, <sup>3</sup>Infection or non-infection diseases, <sup>4</sup>Injury by boat accident, n = number of turtles

The average and range of size for each species are shown in Table 2. The life history stages are shown in Table 2, most stranded sea turtles were juveniles (65.38%, n = 221), followed by sub-adults (20.12%, n = 68) and adults (14.50%, n = 49). Of 149 turtles whose sex could be determined, 120 were female (80.5%).

**Table 2.** The size of stranded sea turtles in the Central Gulf of Thailand

| Turtle | Long |  | Wide |  | Weight |  |
| --- | --- | --- | --- | --- | --- | --- |
|  | Range | Average | Range | Average | Range | Average |
| | (cm) | (mean ± SD) | (cm) | (mean ± SD) | (kg) | (mean ± SD) |
| Green sea turtles | 15–205 | 57.38± 11.13 | 10–98 | 51.20± 9.40 | 0.3–103 | 24.38 ± 13.89 |
| Hawksbill sea turtles | 5.5–90 | 55.09± 14.34 | 5–85 | 49.61± 12.07 | 0.2–90 | 22.84 ± 15.23 |
| Olive ridley sea turtles | 33–90 | 57.66 ± 13.71 | 36–69 | 51.87 ± 13.71 | 15–50 | 25.19 ± 17.25 |
| Leatherback sea turtles | 30–83 | 49.45 ± 14.14 | 40–76 | 45.01 ± 12.65 | 8–53 | 16.71 ± 14.39 |

### 3.2. Causes of stranding

The possible causes of stranding (Table 1) were connected to macroplastics, health (infection) and injury by boat accident, and the unknown cause. From 203 stranded turtles which a cause could be determined, the macroplastics (entanglement or ingestion) were the leading cause (74. 88%, n = 152) followed by health issues (infection and non-infection diseases) (19.70%, n = 40) or injuries from boat accidents (5.42%, n = 11).

Association between marine macroplastics and stranded sea turtles (n = 152) was shown in Table 3, green sea turtles were the most stranded sea turtles to be found with macroplastics (71.05%, n = 108), followed by hawksbill sea turtles (27.63%, n = 42), while olive ridley sea turtle were rare (1.32%, n = 2), and the macroplastic was not found with leatherback turtles. Notably, the frequency of macroplastics problems varied significantly among sea turtles. (chi-square 12.97, df = 2, p = 0.001). Entanglement (69.74%, n = 106) by plastic was found significantly higher (chi-square 137.44, df = 2, p < 0.00001) than ingestion (18.42%, n = 28) and both (entanglement and ingestion at the same time, 11. 84%, n = 18) (Table 3). When compare between two key species, green sea turtles and hawkbills sea turtle, hawksbill sea turtles (90.48%, n = 38/42) were entangled by macroplastics more frequently (chi-square 12.26, df = 1, p = 0.000462) than in green sea turtles (62.11%, n = 66/108). For plastic ingestion, the percentage of green sea turtles (22.22%, n = 24/108) was significantly higher (chi-square 4.09, df = 1, p = 0.043) than hawksbill turtles (9.52%, n = 4/42) (Table 3).

**Table 3.** Association between marine macroplastics and the viability of stranded sea turtles in the Central Gulf of Thailand.

| Species | Macroplastics causes of stranding<br>(n) |  |  |  |  |  |  |  |  | Total<br>(n) |  |
| --- | --- | --- | --- | --- | --- | --- | --- | --- | --- | --- | --- |
|  | %Entanglement |  |  | % Ingestion |  |  | % Both |  |  | %Live | %Dead |
|  | Live | Dead | Total | Live | Dead | Total | Live | Dead | Total |  |  |
| Green sea turtles<br>(n =108) | 34.26%<br>(37) | 26.85%<br>(29) | 62.11%<br>(66) | 2.78%<br>(3) | 19.44%<br>(21) | 22.22%<br>(24) | 3.70%<br>(4) | 12.97%<br>(14) | 16.67%<br>(18) | 40.74%<br>(44) | 59.26%<br>(64) |
| Hawksbill sea turtles<br>(n = 42) | 73.81%<br>(31) | 16.67%<br>(7) | 90.48%<br>(38) | 0.00%<br>(0) | 9.52%<br>(4) | 9.52%<br>(4) | 0.00%<br>(0) | 0.00%<br>(0) | 0.00%<br>(0) | 73.81%<br>(31) | 26.19%<br>(11) |
| Olive ridley sea turtle<br>(n = 2) | 50.00%<br>(1) | 50.00%<br>(1) | 100.00%<br>(2) | 0.00%<br>(0) | 0.00%<br>(0) | 0.00%<br>(0) | 0.00%<br>(0) | 0.00%<br>(0) | 0.00%<br>(0) | 50.00%<br>(1) | 50.00%<br>(1) |
| **Total number**<br>**(n = 152)** | **45.40%**<br>**(69)** | **24.34%**<br>**(37)** | **69.74%**<br>**(106)** | **1.97%**<br>**(3)** | **16.45%**<br>**(25)** | **18.42%**<br>**(28)** | **2.63%**<br>**(4)** | **9.21%**<br>**(14)** | **11.84%**<br>**(18)** | **50.00%**<br>**(76)** | **50.00%**<br>**(76)** |

### 3.3. Association of macroplastics and the death of sea turtles

The association of macroplastics and the viability of turtles are shown in Table 3. Half of the turtles discovered stranded with macroplastics were dead, and the proportion of dead green sea turtles (59. 26%) were significantly higher than hawksbill sea turtles (chi-square 48.52, df = 2, p < 0.001). Interestingly, the majority of sea turtles entangled in macroplastics survived (65.09%, n = 69/106), whereas most turtles that ingested plastic were dead (89.29%, n = 25/ 28). The survival rate of stranded turtles with plastic entanglement was significantly higher (chi-square 26.34, df = 1, p < 0.00001) than plastic ingestion. Some turtles ingested plastics and survived, and the macroplastic was excreted with faeces (n=7) during rehabilitation at the rescue centre.

Necropsy examinations of all deceased turtles (n = 208) revealed that macroplastics were detected in the gastrointestinal tracts of 46 deceased sea turtles (22.11%) (Table 3). All turtles that ingested plastic showed signs of obstruction of the GI tract (stomach and small intestine) caused by a large mass of macroplastics. Among four species, only green sea turtles and hawksbill sea turtles consumed macroplastics.

### 3.4. Types of macroplastics found in GI tracts and on the beaches

The type of macroplastics by macroscopic and microscopic examination is shown in Table 4 and Fig. 2. The macroplastics found in the GI tract were classified as fishing fibre, bags, foams, straw and hard plastics. The presence of plastic fibre in the turtles’ GI tract (62.30%n = 152/244) was significantly higher than other types of plastics (chi-square 426.09, df = 4, p < 0.00001). Plastic fibre was also found on the beaches more frequently (64. 35%, n = 74/115) than others (chi-square 132.76, df = 3, p < 0.00001).

**Table 4.**
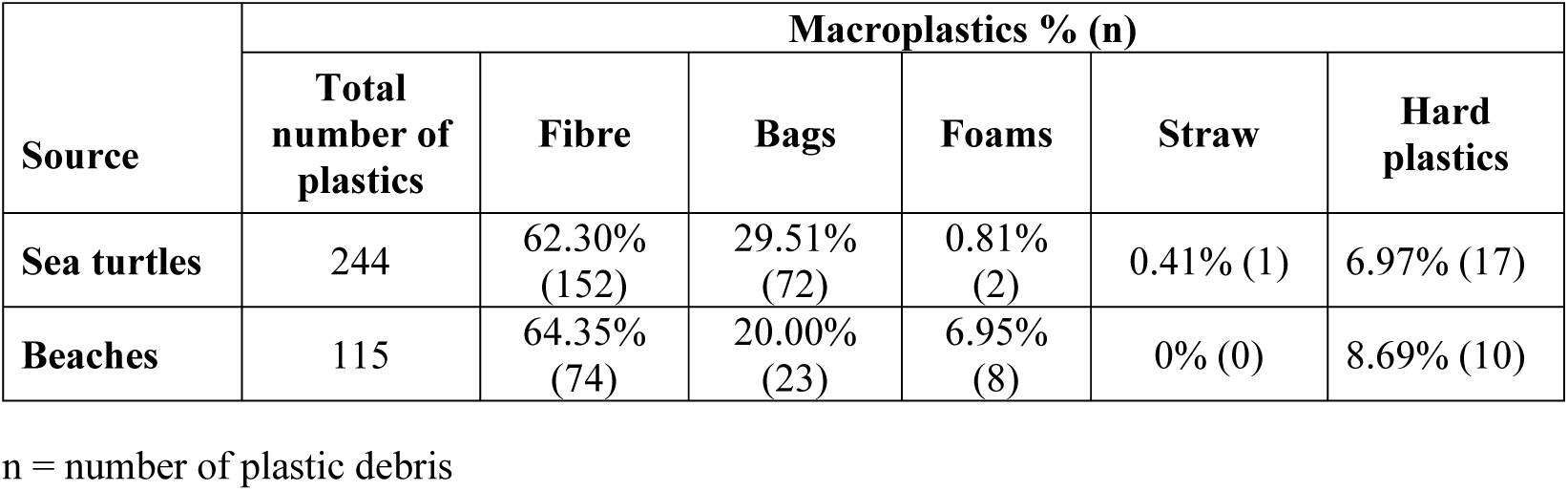
Type of plastic found in the gastrointestinal tract of 30 dead turtles and on the 13 beaches.

| Source | Macroplastics % (n) |  |  |  |  |  |
| --- | --- | --- | --- | --- | --- | --- |
|  | Total number of plastics | Fibre | Bags | Foams | Straw | Hard plastics |
| Sea turtles | 244 | 62.30%<br>(152) | 29.51%<br>(72) | 0.81%<br>(2) | 0.41% (1) | 6.97% (17) |
| Beaches | 115 | 64.35%<br>(74) | 20.00%<br>(23) | 6.95%<br>(8) | 0% (0) | 8.69% (10) |

n = number of plastic debris

**Fig. 2.**
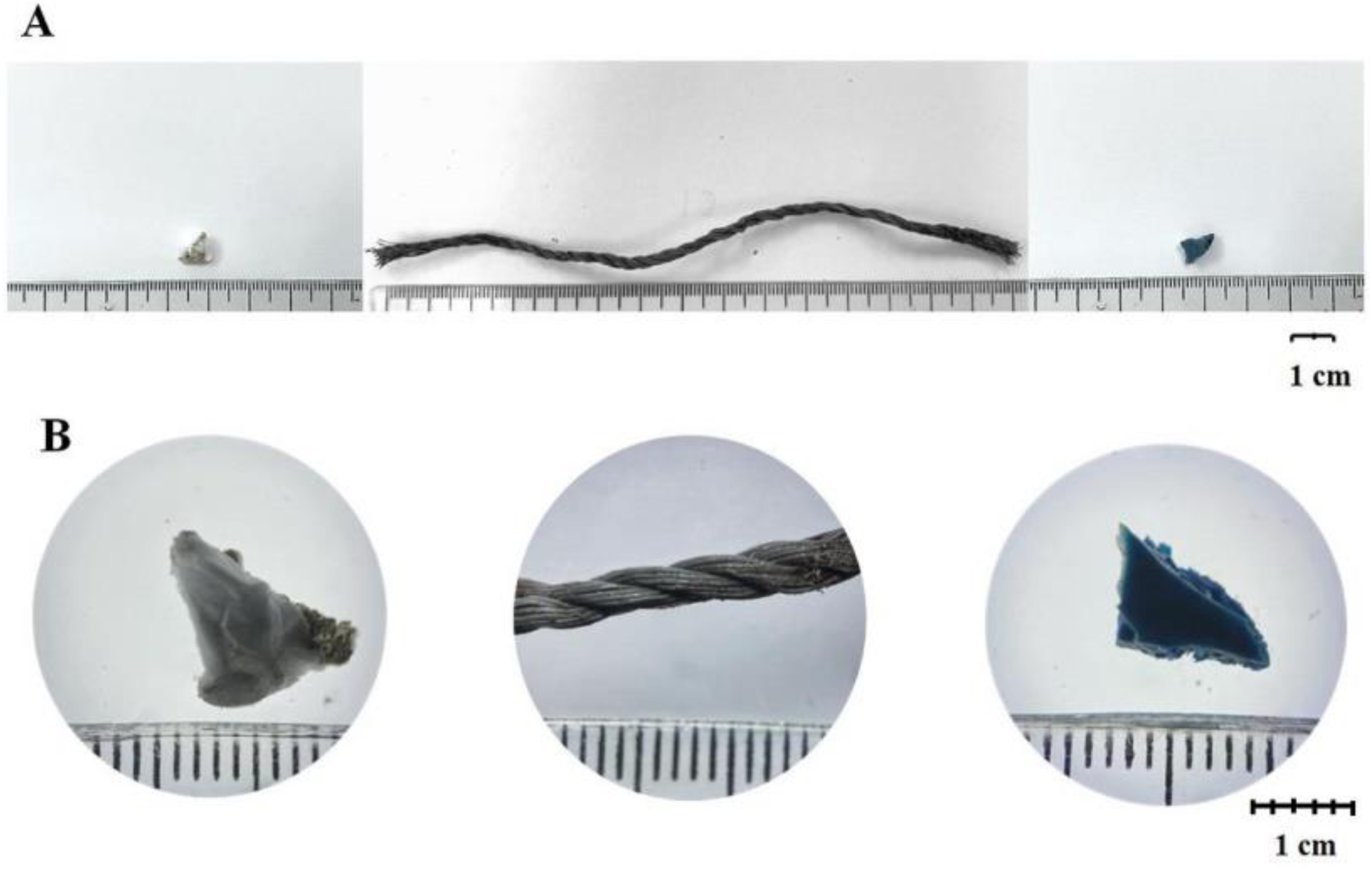
Plastic debris macroscopic and microscopic (A) characteristics of plastic debris and (B) microscopic structure of plastic debris identified visually under a stereo microscope. The left image was foam, the middle image was fibre identified as part of a fishing net, and the right image was hard plastic.

The FTIR derived signature of each plastic found in this experiment was shown Fig. 3. The composition analysis revealed that macroplastics in the GI tract (n = 244) were made from polyethylene (PE) 46.72% (n = 114); polypropylene (PP) 40.98% (n =100), PE+PP 0.82% (n = 2), or nylon 11.48% (n = 28) (Fig. 4). The macroplastics found on the beach (n = 115) were PE 64.35% (n = 74), PP 17.39% (n = 20), PE+PP 6.96% (n = 8), nylon 8.70% (n =10) or polystyrene (PS) 2.61% (n = 3). The proportion of macroplastics made of PE was statically higher than that other materials from both the GI tract (chi-square 194.53, df = 3, p <0.00001) and the beaches (chi-square 185, df = 4, p < 0.00001).

**Fig. 3.**
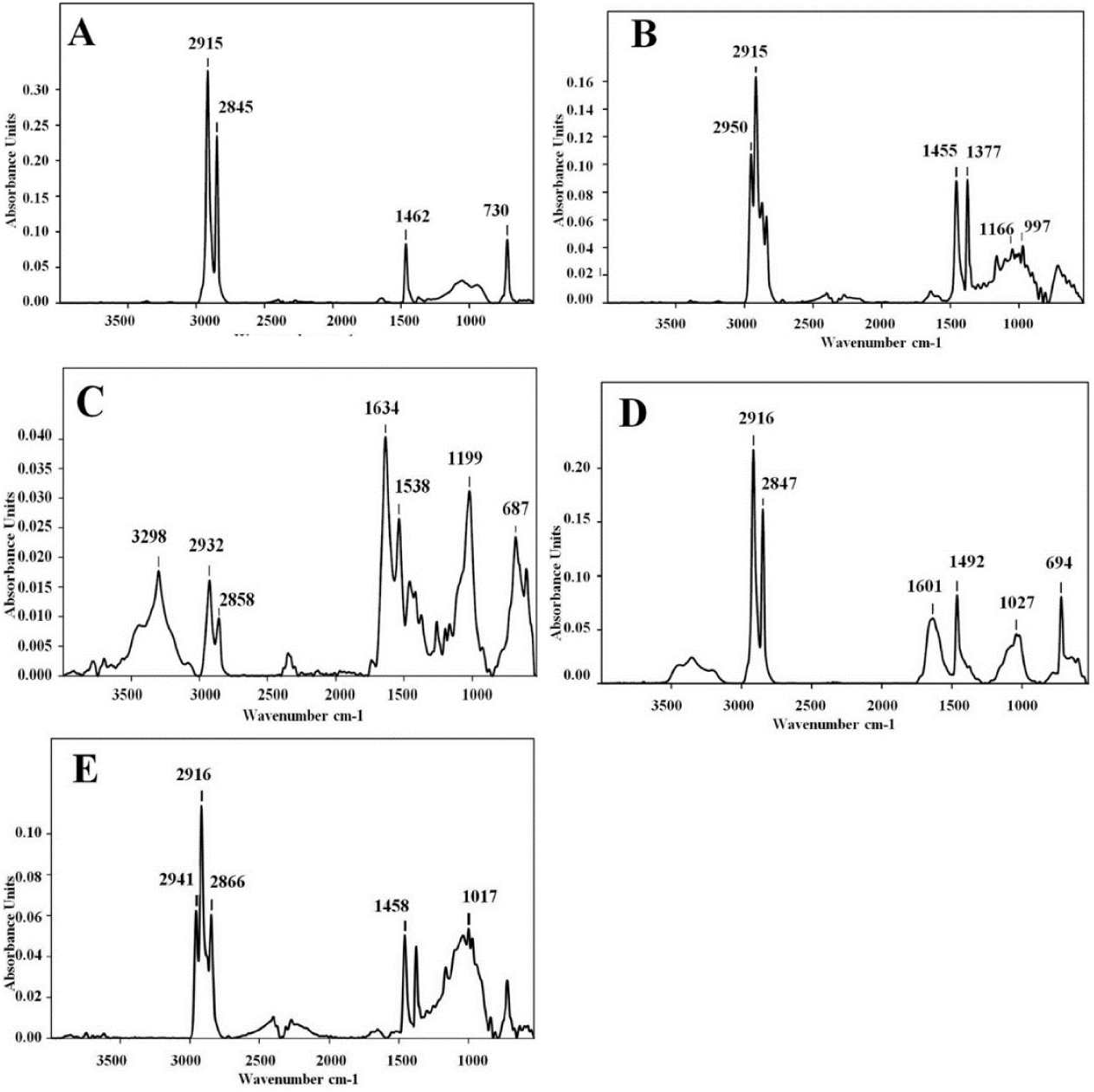
The signature of each macroplastic analysed by FTIR. The characteristic of Polyethylene (A); PE absorbance bands were located at 2915 cm-1, 2845 cm-1, 1462 cm-1 and 730 cm-1, (B) Polypropylene; the characteristic PP absorbance bands were located at 2950 cm-1, 2915 cm-1, 1455 cm-1, 1377 cm-1, 1166 cm-1 and 997 cm-1, (C) Nylon; the characteristic Nylon absorbance bands were located at 3298 cm-1, 2932 cm-1, 2858 cm-1, 1634 cm-1, 1538 cm-1, 1199 cm-1 and 687 cm-1, (D) Polystyrene; the characteristic PS absorbance bands were located at 2916 cm-1, 2847 cm-1, 2858 cm-1, 1601 cm-1, 1492 cm-1, 1027 cm-1 and 694 cm-1 and (E) Copolymer of Polyethylene and Polypropylene (PE+PP); the characteristic Copolymer absorbance bands were located at 2941 cm-1, 2916 cm-1, 2888 cm-1, 1458 cm-1, and 1017 cm-1.

**Fig. 4.**
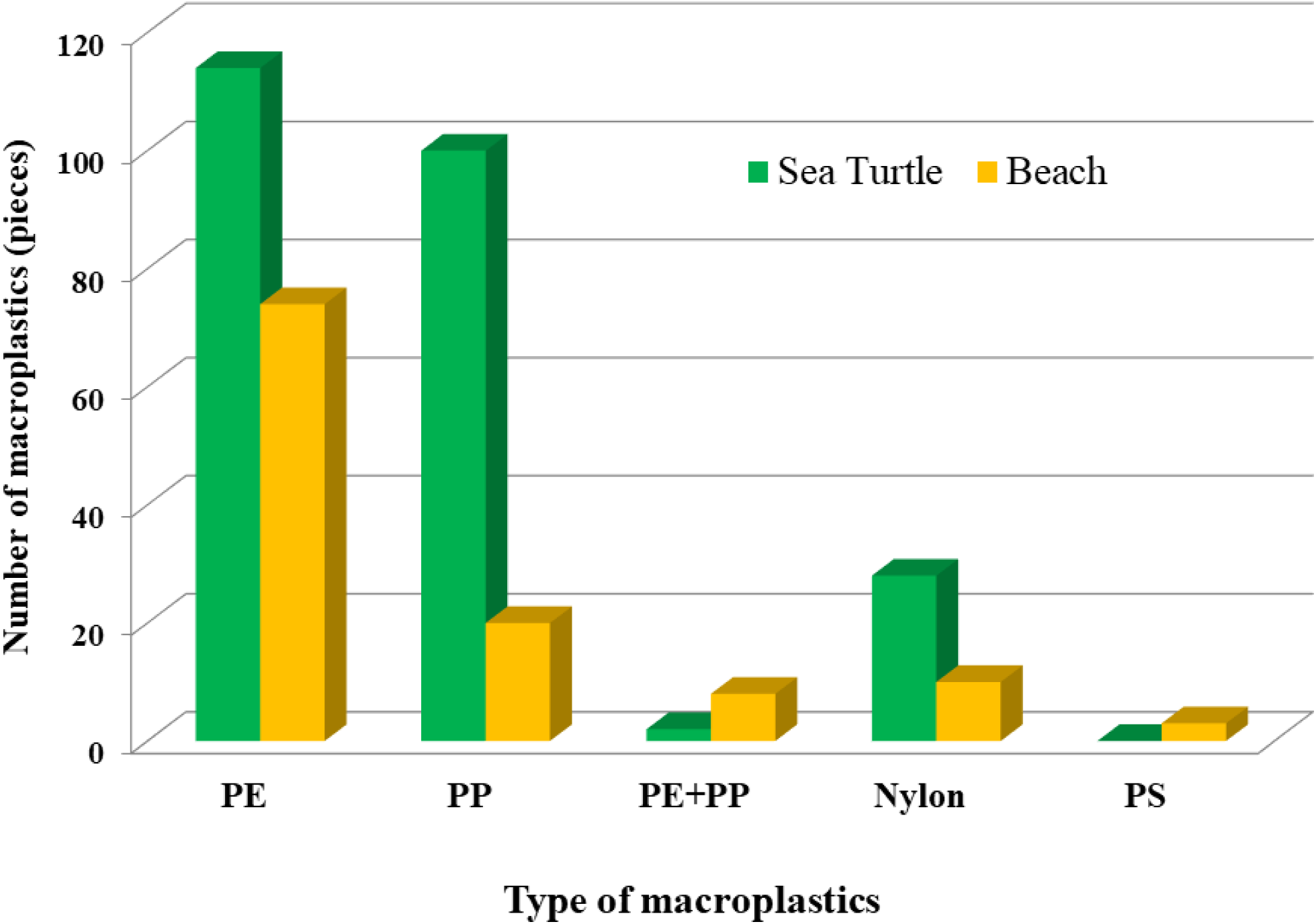
The composition of plastics was determined using FTIR, and samples from the GI tracts of sea turtles and the beach were compared. Polyethylene (PE) and polypropylene (PP) were the most frequently seen plastic waste in turtles and on beaches. Nylon, mixture of PE and PP (PE+PP), and Polystyrene (PS) were rare.

Generally, the fishing fibre can be the fishing net or fishing line. According to FTIR analysis with the reference plastics, fishing nets were made of PP and/or PE, whereas fishing lines were only made of nylon. In the present study, most of the fishing fibres found in the turtles’ GI tract (n =152) were made of PE at 56.58% (n = 86), PP at 26.32% (n = 40), or nylon at 17.11%, (n = 26). Fishing fibres found on the beaches (n = 74) were mainly made of PE at 81.08% (n = 60), PP at 10.81% (n = 8), or nylon at 8.11% (n = 6). Therefore, the majority of plastic fibres found in the GI tract of turtles were fishing nets (82.89%, n = 126/152) similar to fibres found on the beaches (91.89%, n = 68/74), and there was a statistical difference in the type of plastic fibres found in the GI tracts (chi-square 131.57, df = 1, p < 0.00001) and on the beaches (chi-square 103.89, df = 1, p < 0.0001).

### 3.5 Association of macroplastics and life history stages of green sea turtles and hawksbill sea turtles

The association of macroplastics and life history stages of stranded sea turtles are shown in Table 5. Among 3 stages of turtles, only the juvenile stage (n =131) demonstrated a significant correlation with macroplastics (chi-square 36.03, df = 1, p < 0.001). Analysis in both species, there was a statistical difference between entanglement and ingestion in the juvenile stage (chi-square 76.50, df = 1, p < .00001) and the sub-adult stage (chi-square 21.33. p < 0.00001), the juvenile and sub-adult stages had the higher rate of entanglement than ingestion. The percentage of entanglement in adult turtles was significantly lower than in the juvenile stage (chi-square statistic is 7.20, df =1 p = 0.0072) and sub-adult (chi-square 4.44, df = 1, p = 0.035).

**Table 5.** Association of macroplastics to life history stage of major species of stranded sea turtle (green sea and hawksbill) in the Central Gulf of Thailand.

| Species | Stage | Problems with macroplastics (n) |  |  | The presence of macroplastics with turtles (n) |  |
| --- | --- | --- | --- | --- | --- | --- |
|  |  | %Entanglement | %Ingestion | % Both | % Yes | % No |
| **Green sea turtles**<br>**(n = 143)** | Juvenile<br>(n = 82) | 64.62% (42) | 20.00% (13) | 18.38% (10) | 79.7% (65) | 20.73% (17) |
|  | Sub-adult<br>(n = 29) | 65.22% (15) | 13.04% (3) | 21.74% (5) | 79.31% (23) | 20.69% (6) |
|  | Adult<br>(n = 32) | 45.00% (9) | 40.00% (8) | 15.00% (3) | 62.5% (20) | 37.5% (12) |
| **Hawksbill sea turtles**<br>**(n = 58)** | Juvenile<br>(n = 49) | 91.66% (33) | 8.33% (3) | 0.00% (0) | 73.47% (36) | 26.53% (13) |
|  | Sub-adult<br>(n = 9) | 83.33% (5) | 16.67% (1) | 0.00% (0) | 66.67% (6) | 33.33% (3) |
|  | Adult<br>(n = 0) | 0.00% (0) | 0.00% (0) | 0.00% (0) | 0.00% (0) | 00.00% (0) |
| **Both species**<br>**(n =201)** | Juvenile<br>(n =131) | 74.26% (75) | 15.84% (16) | 9.90% (10) | 77.1% (101) | 22.90% (30) |
|  | Sub-adult<br>(n = 38) | 68.97% (20) | 13.79% (4) | 17.24% (5) | 76.20% (29) | 23.68% (9) |
|  | Adult<br>(n = 32) | 45.00% (9) | 40.00% (8) | 15.00% (3) | 62.50% (20) | 37.50% (12) |

## 4. DISCUSSION

In the present study, the green sea turtle was the most frequently stranded sea turtle in the Central Gulf of Thailand, accounting for approximately 74 % of the total, followed by the hawksbill sea turtle accounted for approximately 21%. The high rate of stranding in green sea turtles can be simply explained by the fact that green sea turtles and hawksbill sea turtles are the first and second significant populations, respectively, whereas olive ridley sea turtles and leatherback sea turtles were rare in the Gulf of Thailand according to the report of Marine and Coastal Resources Research and Development Institute (https://dmcrth.dmcr.go.th).

The incidence of macroplastic ingestion in the present study was highest in green sea turtles, followed by hawksbill sea turtles. In comparison between the two main species, entanglement was more prevalent in hawksbill sea turtles than green sea turtles, whereas a greater proportion of green sea turtles consumed macroplastics. This phenomenon can be explained by the feeding behaviour of these two species. Green sea turtles are primarily herbivores, usually consuming seagrass and seaweed (Carrión-Cortez et al. 2010, Awabdi et al. 2013, Stokes et al. 2019), of which the morphology is similar to the remnants of fishing nets/lines. Additionally, ghost nets and fishing lines are more easily entangled on a rocky substratum where seaweed grows, and this may cause turtles to accidentally consume plastics (Thiel et al. 2018). In contrast, hawksbill sea turtles feed primarily on small animals such as fish, sponges, and algae (Meylan 1988, Leon & Bjorndal, 2002, Bell 2012). This plastic ingestion behaviour in this study was similar to that observed in other studies conducted in the Brazilian ocean (Carman et al. 2014, Rizzie et al. 2019). Due to the high incidence of entanglement rate in hawksbill turtles, it is possible that they were attempting to capture small marine animals already entangled in the net. It has been shown that Thailand’s ghost net is capable of trapping a large number of small marine animals (Sukhsangchan et al. 2020, Ballesteros et al. 2018).

A related research conducted on the eastern coast of the United Arab Emirates discovered that the majority of plastics found in the gastrointestinal tracts of green sea turtles is the plastic fibre (Yaghmour et al. 2018). Conversely, studies in the Pacific Ocean near Hawaii found that most of plastics found in the GI tract of green sea turtles are fragments of hard plastics, while the plastic fibre is rare (Wedemeyer-Strombel et al. 2015, Clukey et al. 2017). Notably, most macroplastics found in the present study were made of PE, a material similar to that found on the beaches, implying that turtles may consume the most abundant macroplastics in the Central Gulf of Thailand. These macroplastics that entangled turtles or being ingested are probably located close to the shore, as data from the Marine and Coastal Resources Research Centre revealed that sea turtles in the Gulf of Thailand rarely venture from the shore (https://km.dmcr.go.th/en/c_6/d_982). As a result, we believed that the type of plastic found in the gastrointestinal tracts of stranded sea turtles in each location represented the most abundant macroplastics found in the ocean or on the shore.

In the Thai ocean, olive ridley sea turtles and leatherback sea turtles are rare (https://dmcrth.dmcr.go.th). They accounted for approximately 4% of stranded turtles discovered in this study, and the primary cause of stranding was unknown (85%). Only 18% of olive ridley sea turtles were found entangled with macroplastics, while the standing of leatherback sea turtles was not associated with macroplastics. The study in the Southern Ocean of Brazil found a similar finding to ours; olive ridley sea turtles have the lowest plastic ingestion rate among sea turtles (Rizzie et al., 2019). Conversely, the study in other areas has discovered a high incidence (up to 100%) of olive ridley sea turtles ingesting plastics (Wedemeyer-Strombel et al. 2015, Clukey et al. 2017, Jung et al. 2018b). For the leatherback sea turtles, the occurrence of plastic ingestion is relatively low; one study reveals no incidence (Clukey et al. 2017), and another reports 34% (Mrosovsky et al. 2009). The most common form of plastic debris found in the GI tract of leatherback sea turtles is bags which may imitate the main diet of leatherback sea turtles, gelatinous animals such as jellyfish (Mrosovsky et al. 2009, Heaslip et al. 2012).

The majority of known causes for the stranding of sea turtles in the Central Gulf of Thailand were entanglement by fishing nets (also known as ghost nets). This is unsurprising, given Thailand’s position as a leading exporter of edible fisheries products and the Central Gulf of Thailand’s importance as a main area for fisheries according to data from Food and Agriculture Organisation of The United Nations (http://www.fao.org/fishery/facp/THA/en). Ghost nets are a global issue because more than half-million tons of fishing nets are lost and discarded annually, causing the entanglement of large marine animals, especially sea turtles (Wilcox et al. 2013). Some ghost nets can be tracked back to their country of origin, with the majority coming from Asian countries such as Thailand (Gunn et al. 2010). Another risk analysis study indicated that South-East Asia, particularly Thailand, might be one of the highest risk places for sea turtles to ingest the plastics or become entangled (Schuyler et al. 2016, Duncan et al. 2017). The elimination of ghost nets in the Gulf of Thailand should be the first priority for resolving the turtle stranding crisis. However, no national legislation regulating ghost nets has been established yet. Only few activities by government agencies, non-profit organizations, and local people to clear ghost nets from the ocean were reported by the media (https://www.diveagainstdebris.org/).

We discovered that most stranded sea turtles were juvenile, and the majority of them were associated with macroplastics, causing the entanglement more than ingestion. This finding is consistent with a previous study indicating that juvenile green sea turtles and hawksbill turtles are more susceptible to entanglement than other stages (Duncan et al. 2017). Entangled animals may drown, particularly smaller animals may drown immediately if the fishing gear is large or heavy. They also starve and may suffer physical trauma and infections from the fishing gear cut into their flesh. (https://www.fisheries.noaa.gov/insight/entanglement-marine-life-risks-and-response). The ingestion of plastic debris of sea turtles begins since hatching (Eastman et al. 2020); juvenile turtles face a greater risk of death from plastic ingestion than adults because they consume a greater quantity of plastic than adults do despite their smaller bodies (Wilcox et al. 2018). Consuming an excessive amount of macroplastics can obstruct the gastrointestinal tract and result in acute death (Stamper et al. 2009), while a small amount of plastic can have sub-lethal effects through increasing satiety, inhibiting digestion, and impairing absorption, resulting in malnutrition and weakness (Santos et al. 2020). Notably, both conventional and biodegradable plastics are toxic to turtles due to their difficulty degrading by digestive juice (Müller et al. 2012).

In conclusion, this study proposes that macroplastics, mainly fishing nets made of PE and PP, constitute a significant cause of the stranding of sea turtles in the Central Gulf of Thailand. These macroplastics may cause weakness, unwellness, and eventually death in sea turtles, especially the green sea turtles and hawksbill sea turtles. Therefore, to prevent more sea turtle deaths in the Central Gulf of Thailand, it is critical to urgently reduce and eliminate marine plastic debris particularly fishing nets.

## Acknowledgements

This project was supported by Walailak University’s Personal Research Grant (WU-IRG-63-033), the New Strategic Research project (P2P 2564 Phase 1), and the Marine Animal Research and Rescue Centre of Walailak University. We would like to Thank the English proofreading service from Research Unit of Akkhraratchakumari Veterinary College.

